# Fishery Cooperatives as Institutional Intermediaries in Fragile Contexts: Assessing Social-Ecological Resilience in South-Central Somalia

**DOI:** 10.64898/2026.07.25.740677

**Authors:** Anas A. Madar, Omar H. Hadaaf

## Abstract

Small-scale fisheries provide essential coastal livelihoods in developing regions, yet they often confront severe institutional voids within fragile-state contexts. Adopting a social-ecological systems framework, this study addresses three sequential objectives: characterizing the contemporary institutional and demographic status of fishery cooperatives across the five South-Central Federal Member States in comparison to their late-1980s historical peak; ranking the socio-economic, infrastructural, regulatory, and environmental impediments to cooperative efficacy; and evaluating the contributions of these cooperatives to sustainable fisheries development. This investigation analyzes how fishery cooperatives act as institutional intermediaries to bolster economic resilience within the Somali coastal economy—a sector central to the National Transformation Plan, despite persistent regulatory fragmentation. Through semi-structured qualitative interviews with key informants—analyzed via reflexive thematic analysis and chi-square testing of age-cohort structure—the study finds that illegal, unreported, and unregulated fishing and inadequate post-harvest infrastructure are the principal constraints, whereas training, education, and community collaboration constitute the primary contributions. The results characterize these cooperatives as hybrid institutions that compensate for limited central-state capacity in remote coastal settlements while serving as intermediaries for the Federal Ministry of Fisheries and Blue Economy where co-management mandates are implemented. By extending the actor-community dimension of the SES framework to include age-cohort structure, this research demonstrates that clan-based reciprocal arrangements can serve as functional substitutes for Ostrom’s first design principle in contexts of persistent fragility. The study recommends multi-stakeholder partnerships to advance targeted capacity-building, financial literacy, and technological integration, and suggests future research into the post-harvest and retail sectors to examine the role of women in the Somali blue economy.

## 1. Introduction

Coastal populations in developing regions depend heavily on small-scale fisheries for both subsistence and economic stability. This reliance is increasingly threatened by climate change, deficient institutional oversight, and the intrusion of foreign fishing fleets (Turakulov et al., 2025). Although scholars frequently frame harvester cooperatives as critical instruments for collective action that mitigate externalities and stabilize open-access resource systems, such literature has largely emerged from stable political contexts rather than fragile states. This study seeks to address two primary empirical gaps. First, the institutional determinants of cooperative resilience within fragile federations remain under-theorized regarding the social-ecological systems framework. Second, the potential for customary clan governance to act as a functional substitute for Ostrom’s first design principle—a mechanism often examined within Latin American settings—has not been sufficiently investigated in fishery co-management. Somalia offers a unique venue to address these lacunae, given its significant ecological and political profile. The nation possesses Africa’s second-longest coastline, extending approximately 3,025 km along the Gulf of Aden and the Indian Ocean, alongside an Exclusive Economic Zone of roughly 825,000 km² in the Western Indian Ocean (Breuil & Grima, 2014; Kaba, 2024). Furthermore, the federalized fisheries governance framework implemented since 2017 situates South-Central cooperatives at the intersection of state authority, traditional clan governance, and the FAO Voluntary Guidelines for Securing Sustainable Small-Scale Fisheries in the Context of Food Security and Poverty Eradication (Nakamura et al., 2021).

Somalia’s National Transformation Plan emphasizes the fisheries sector as central to inclusive economic growth, aligning with Sustainable Development Goal 14.b regarding equitable access for small-scale artisanal fishers to resources and markets. This agenda is enacted through the Somali Fisheries Development Framework 2018–2020 under the Eighth National Development Plan (Mohamed, 2022), applying co-management strategies consistent with established Voluntary Guidelines (Nakamura et al., 2021) under the coordination of the Federal Ministry of Fisheries and Blue Economy. Within this structure, fishery cooperatives serve as vital community-level institutional interfaces. Despite this policy architecture, the sector remains largely “unregulated and uncontrolled” (Hassan & Hossain, 2023; Sumaila & Bawumia, 2014), sustaining significant livelihood vulnerabilities notwithstanding the availability of productive fishing grounds in the Gulf of Aden and the Indo-Pacific.

We argue that South-Central Somali fisheries cooperatives occupy a hybrid position on a state-engagement continuum, functioning simultaneously as substitutes for absent central-state capacity in remote settlements and as institutional intermediaries for the Federal Ministry of Fisheries and Blue Economy where post-2017 regulatory frameworks are active. The balance between these roles varies across the five Federal Member States, shaped by the local penetration of the Federal Blue Economy Framework (Abdalla, 2024), the prevalence of customary clan governance (ABDIRAHMAN, 2022), and the cooperative’s registration status (Abdalla, 2024). While Somali cooperatives operate without Ostrom’s first design principle, they utilize informal clan-based governance as a functional substitute—a mechanism noted in broader Somali co-management literature (ABDIRAHMAN, 2022) but previously untheorized for the South-Central region. This study addresses this gap by applying the social-ecological systems framework at the member-state level and incorporating age-cohort structure as a determinant of governance preferences.

### 1.2 Primary Research Question

To what extent do fishery cooperatives in the five South-Central Federal Member States of Somalia function as institutional intermediaries within the social-ecological system, and which factors account for the variation in their contributions to institutional resilience across these jurisdictions?

#### 1.2.1 Specific Research Questions

1. What constitutes the contemporary institutional and demographic profile of fishery cooperatives across the five South-Central Federal Member States?
2. What are the primary impediments constraining cooperative efficacy within these states, and how are these constraints hierarchically ranked by prevalence?
3. Through which primary mechanisms do fishery cooperatives most significantly facilitate sustainable fisheries development in the South-Central Federal Member States?

## 2. Literature review and theoretical framework

### 2.1 Theoretical framework

The resilience and sustainability of South-Central Somalia’s coastal fisheries depend on the institutional alignment of the four first-tier Social-Ecological Systems variables: resource systems, resource units, governance systems, and actor communities. This institutional alignment, as operationalized by Somali fishery cooperatives, is a key determinant of successful management. However, these efforts are increasingly challenged by the persistent impact of Illegal, Unreported, and Unregulated fishing.

### 2.2 Concept of fishery cooperatives in Somalia

The formal institutional framework for Somali fishery cooperatives began with Law No. 40 of 1973, which established them as “voluntary associations of individuals with common economic interests, organized on a democratic basis” (Yassin, 1981). Promulgated under the Siad Barre administration’s socialist economic agenda, the Cooperatives Development Act received support from Soviet-aligned development programs across various sectors (Roberts et al., 2019). By the late 1970s, the Ministry of Fisheries and Marine Resources had registered 18 cooperatives along the coast from Ras Kambone to Zaylac (Yassin, 1981). Additionally, the 1975 Dabadheir famine prompted the government to resettle approximately 15,000 displaced pastoralists in settlements such as Brawe, El-Ahmed, Eyl, and Adale, incorporating them into fishing cooperatives to diversify livelihoods (Yassin, 1981). This model reached its peak in 1988, comprising 65 cooperatives and roughly 3,500 members (Perry & Aden, 2023). Following the 1991 collapse of the central state, however, most cooperatives dissolved due to the absence of a supervisory authority (Glaser et al., 2019).

*The Somali Federal Ministry of Fisheries and Marine Resources has reactivated the policy framework — not the legal one — through the Somali Fisheries Development Framework 2018–2020 under NDP8* (Mohamed, 2022)*, the 8th-to-9th NDP transition with NDP9 reserving ‘large portions of initiating Somalia’s blue economy sectors’* (Mohamed, 2022)*, and the IGAD-validated National Baseline Assessment Report of the Blue Economy’s contribution to sustainable economic development* (Mohamed, 2022)*; however, the underlying fisheries legal framework remains largely unrestored, with most investment regulations dated before 1991, no organizational statute covering offshore permitting, leasing, and monitoring, and the laws needed to operationalize the provisional constitution’s independent judiciary still absent* (Mohamed, 2022). *Today, 20 fishery cooperatives are federally registered with the Federal Ministry’s public registry, while an additional layer of clan-customary autonomous cooperatives operates in sub-state coastal districts without formal registration, reflecting the substitution-mode extreme of the cooperative continuum anticipated in the Voluntary Guidelines for SSF*(ABDIRAHMAN, 2022).

*Fisheries governance scholarship conceptualizes the cooperative model as a collective-action institution for the management of common-pool resources* (Elsler et al., 2022). Leveraging Ostrom’s eight design principles for successful common-pool resource institutions—as reformulated within the social-ecological systems framework (McGinnis & Остром, 2014)—this study examines Somali fishery cooperatives as institutional intermediaries. Specifically, their internal governance structures are evaluated for functional adherence to principles 1, 3, 4, and 7 within the context of South-Central Somalia (Elsler et al., 2022). Given Somalia’s extensive coastline—spanning approximately 3,025 kilometers along the Gulf of Aden and the Indian Ocean, ranking among the longest in continental Africa (Glaser et al., 2015)—the cooperative model assumes particular academic significance. This relevance is underscored by the sector’s largely unregulated status; in the prevailing post-conflict institutional vacuum, any form of collective governance represents an analytically valuable subject of study (Roberts et al., 2019).

### 2.3 Challenges of Somali Fishery Cooperatives

The literature identifies four primary constraints to the efficacy of Somali fishery cooperatives. First, institutional capacity remains severely limited by rudimentary governance frameworks, insufficient financial management expertise, and a lack of technical training for both leadership and membership (ABDIRAHMAN, 2022). Second, technical impediments—including antiquated fishing gear and traditional, low-efficiency harvesting methods—substantially diminish potential yield and resource utilization (ABDIRAHMAN, 2022). Third, systemic infrastructure deficiencies, particularly the absence of cold-chain and post-harvest facilities, restrict cooperatives to lower-value domestic trade, thereby precluding entry into high-value export markets (Mohamed, 2022). Finally, environmental and resource-related stressors, such as climate-induced fluctuations in fish populations and recurrent extreme weather, destabilize both coastal infrastructure and artisanal fishing operations, contributing to broader socioeconomic volatility (Glaser et al., 2019; Sumaila & Bawumia, 2014).

These constraints create a significant gap between current practices and the governance structures recommended by Ostrom’s principles for sustainable resource management. Specifically, institutional weaknesses hinder adherence to Principle 3, while technical and resource limitations restrict the monitoring needed for Principle 4 (ABDIRAHMAN, 2022). This issue is evident in Mogadishu, where the Somali Coast Guard and regional fisheries administrators face severe capacity constraints—such as limited technical expertise, insufficient personnel, and a lack of systematic data—that impede regulatory compliance (ABDIRAHMAN, 2022). Furthermore, reliance on sporadic, situational sanctioning reveals a broader systemic failure to ensure consistent institutional enforcement (ABDIRAHMAN, 2022).

### 2.4 Somalia Domestic Fish Catch

*The Somali domestic marine catch in 2006 totaled 32,600 tonnes. Relative to a Western Indian Ocean Exclusive Economic Zone of approximately 830,400 km² characterized by a primary productivity of roughly 882 mg C/m²/day, this figure indicates that current production accounts for less than 6% of the estimated 600,000-tonne annual bio-ecological potential* (Hassan & Hossain, 2023). The significant disparity between realized output and latent capacity, when situated within the broader fisheries development challenges characteristic of fragile states (Sumaila & Bawumia, 2014), highlights the imperative to prioritize cooperative production capacity as a national objective (Roberts et al., 2019). Consequently, long-term sector development necessitates targeted investment in infrastructure, capacity building for artisanal fishers, and the institutional strengthening of cooperative organizations (Glaser et al., 2015).

### 2.5 Contribution of Cooperatives to Sustainable

#### 2.5.1 Fisheries Development

Fishery cooperatives across the African continent serve as critical institutional mechanisms for fostering sustainable development and economic resilience within coastal populations (Asante, 2024). The scholarly literature is underpinned by four prominent regional analogues. In Kenya, Beach Management Units have facilitated resource co-management and enhanced regulatory compliance. South African cooperatives have empowered small-scale fishers through the provision of collective fishing rights and vocational training. Similarly, Ghanaian canoe-fishers’ associations have utilized collective bargaining to regulate market pricing and mitigate illegal fishing practices. Finally, Tanzania’s BMUs—comprising 179 units established under the Fisheries Act No. 22 of 2003 and the Fisheries Regulations of 2009—demonstrate the efficacy of institutionalized community stewardship. Within this framework, 68 units possess formal management plans, and 39 operate under legally enforceable District Council by-laws.

Furthermore, higher-tier Collaborative Fisheries Management Areas in the Rufiji, Mafia, and Kilwa districts have successfully operationalized marine stewardship through localized gear and area restrictions, alongside the documented integration of women in gleaning and seaweed-cultivation activities (Elegbede et al., 2023). These regional analogues suggest five primary contribution pathways relevant to Somali fishery cooperatives. First, training and education serve as the cornerstone for translating collective-action governance into positive social-ecological outcomes. Capacity building in financial management and transparent governance is highly recommended in the Ghanaian context (Asante, 2024), while robust collective action—characterized by collective-choice agreements, active monitoring, conflict-resolution mechanisms, and nested enterprise arrangements—is quantitatively linked to the sustainability of cooperative fisheries at the national level (Elsler et al., 2022). Second, community collaboration and technology promotion facilitate the dissemination of catch and post-harvest technologies among members (Glaser et al., 2019). Third, policy advocacy and governance representation ensure the inclusion of fisher perspectives within central policy cycles (Roberts et al., 2019a, 2019b). Fourth, market access and fair pricing represent the primary direct economic benefit for members operating within volatile fish markets (Perry & Aden, 2023). Fifth, credit and financial services provide the financial inclusion necessary for operational scalability and help overcome historical barriers to cooperative development in Somalia (Ibrahim & Ngina, 2019).

These pathways allow cooperatives to consolidate small-scale fishing efforts through specific operational mechanisms. Community collaboration—defined here as the cooperative-mediated pooling of pre-trip procurement, the exchange of beach-landing catch data, and customary clan-based reciprocal labor arrangements—substitutes for Ostrom’s Principle 1 in Somali small-scale fisheries, aligning with social-ecological systems collective-action literature (Elsler et al., 2022) and echoing leadership-trust drivers seen in Kenyan Beach Management Units and Mexican collective-choice agreements (Murunga et al., 2021). Technology promotion is operationalized through AIS-based catch reporting and partnerships in post-harvest solar cold storage (Glaser et al., 2019). Furthermore, policy advocacy, augmented by credit and financial services, enhances catch-monitoring accuracy through co-management mandates under the Federal Blue Economy Framework (Abdalla, 2024). When supported by credit, these market-access and fair-pricing initiatives strengthen the bargaining power of cooperatives to secure equitable trade terms. By integrating these strategies, fishery cooperatives are positioned to leverage Somalia’s estimated EEZ potential—capped at an annual catch of 600,000 tonnes within an 830,400 km² EEZ with a primary productivity of roughly 882 mg C/m²/day (Hassan & Hossain, 2023)—to realize the capacity gains evidenced in this study (Abdalla, 2024).

### 2.6 Gap and Theoretical Contribution

This study addresses four primary gaps in the literature. First, post-conflict sectoral reactivation remains unquantified across the five South-Central Federal Member States. Second, while fisheries-sector challenges are conceptualized, their relative frequencies lack systematic reporting across comparable empirical samples. Third, the five contribution pathways have not been jointly evaluated within these states. Finally, scholarship has yet to investigate the compensatory mechanisms enabling Somali cooperatives to maintain collective-choice arrangements despite the absence of Ostrom’s Principle 1. Specifically, this study analyzes how clan-based informal structures and social capital function as substitutes for the formal boundary-clarification mechanisms delineated in Principle 1 (Roberts et al., 2019).

## 3. Research Methodology

This research uses a qualitative-dominant mixed-methods approach, integrating semi-structured key-informant interviews with a chi-square test of independence to assess a specific demographic hypothesis.

### 3.1 Study Area and Sampling

The study area encompasses the five South-Central Federal Member States adjacent to the Indian Ocean and Gulf of Aden. These states were selected through maximum-variation purposive sampling, based on three dimensions aligned with Social-Ecological System first-tier variables: infrastructure variability, evidenced by documented cold-chain deficiencies and a 60% reliance on small-scale fisheries gillnets, lines, and traps (Breuil & Grima, 2014); governance variability, evaluated via federal-penetration metrics under the Federal Blue Economy Framework and corresponding co-management mandates (Abdalla, 2024); and exposure to Illegal, Unreported, and Unregulated fishing, as quantified by Secure Fisheries AIS data, which estimated an annual foreign-driven IUU catch of 300,000–400,000 metric tons in Somali waters(Glaser et al., 2019). This five-state stratified sampling strategy ensures robust sub-national comparability (ABDIRAHMAN, 2022).

The target population comprises adult members and officials of fishery cooperatives. A multi-stage purposive sampling approach was employed, selecting three federally registered cooperatives per state, yielding a total sample of *n* = 90 key informants. Sampling proceeded until thematic saturation—customarily achieved between the 18th and 22nd informants per state—was reached. Given that 15 interviewees per stratum typically suffice for thematic saturation in homogeneous purposive samples, the quota of 18 informants per state exceeds standard requirements, ensuring the reliability of state-level comparisons.

### 3.2 Data collection and analysis

Semi-structured interviews were conducted between March and May 2026, approximately one hour each at informant-chosen locations. The researchers recorded the interviews with informed consent and transcribed them verbatim. A reflexive thematic analysis following Braun and Clarke’s six-phase procedure — with a directed top-level code scaffold drawn from the SES framework’s four first-tier subsystems (McGinnis & Остром, 2014), followed by inductively derived subordinate codes that converge on themes along the action-situations *→ outcomes arrows — is applied as the interpretive procedure for the 90 key-informant transcripts* (Campbell et al., 2021). Age-cohort differences in attitudes toward policy advocacy and modern governance-technology integration are tested with the χ² test of independence (α = 0.05, df = 1) after verifying the expected-cell-frequency assumption (no expected cell below five).

## 4. Results

This section presents the demographic profile of the study participants. It then ranks the six primary fisheries-sector challenges identified through thematic analysis, summarizes cooperative contribution pathways, and reports the χ² test results for Hypothesis H3.

### 4.1 Sample Demographic Profile

The 90 key informants were stratified equally across the five South-Central Federal Member States, drawn from a population of 65 federally registered fishery cooperatives through the purposive selection of three cooperatives per state.

**Table 1.** Gender Distribution of the Respondents.

| **Gender** | **Frequency** | **Percent** |
| --- | --- | --- |
| Female | 15 | 16.7% |
| Male | 75 | 83.3% |
| **Total** | **90** | **100%** |

The participant sample—comprising 83.3% male and 16.7% female respondents—reflects the established gender-segmented division of labor within South-Central Somali small-scale fisheries. In this sector, men traditionally lead in boat-handling and gear operations, while women are largely concentrated in post-harvest processing and beach-side trading, as observed at Mogadishu’s Liido, Urubo, and Abaydhahn landing sites (Hassan & Hossain, 2023). This male-dominated occupational profile, where female involvement is primarily restricted to fish trading, is reaffirmed by local market data (Perry & Aden, 2023) and aligns with FAO benchmarks, which estimate direct capture participation at approximately 14%, increasing to roughly 50% when including onshore post-harvest activities (Mohamed et al., 2026).

**Table 2.** Age of the Respondents.

| Age Cohort | Frequency | Percent |
| --- | --- | --- |
| 18–25 years | 10 | 11.1% |
| 26–35 years | 33 | 36.7% |
| 36–45 years | 34 | 37.8% |
| 46–55 years | 8 | 8.9% |
| 56+ years | 5 | 5.5% |
| **Total** | **90** | **100%** |

*The age distribution exhibits a right skew toward working-age and mature cohorts, which are documented as the demographically dominant segments in Somali small-scale fisheries. This distribution reflects the educational profile reported for the Mogadishu SSF labor force—characterized by 48% illiteracy and approximately 20% secondary-educated status—correlating with adult, post-formal-education male-occupation cohorts* (Hassan & Hossain, 2023). These findings also align with the demographic profiles for South-Central coastal fishers detailed in the *Baseline Report Somalia* (Breuil & Grima, 2014). Regarding specific cohorts, the 36–45 and 26–35 age groups represent the modal segments, accounting for 37.8% and 36.7% of respondents, respectively, while the 18–25 (11.1%), 46–55 (8.9%), and 56+ (5.5%) cohorts form smaller subsets. Overall, the 85.6% working-age aggregate is consistent with the regional SSF bias toward able-bodied adult fishers noted in prior governance assessments (ABDIRAHMAN, 2022). Furthermore, the binary split of 18–35 versus 36+ used for the H3 chi-square test follows the African Union Youth Charter threshold, a widely accepted metric in ILO and FAO SSF policy tracking for differentiating youth-versus senior-driven technological adoption. This pre-registration is detailed here to ensure the transparency of the test’s inferential framework (Nakamura, Chuenpagdee, & Halimi, 2021).

### 4.2 Age-Cohort Hypothesis Test

To evaluate the association between age cohorts and attitudes toward policy advocacy and the integration of modern governance technologies, a two-tailed *χ*^2^ test of independence was conducted (*α* = 0.05,*n* = 90). As the calculated expected cell frequencies (*E* ≈ 21.02,21.98,22.98,24.02) surpassed the minimum threshold of 5 required for the asymptotic *χ*^2^ distribution, the application of Fisher’s exact test was deemed unnecessary. The analysis yielded *χ*^2^ ≈ 8.68 (*p* ≈ 0.003) with a Cramér’s *V* ≈ 0.31, indicating a medium effect size. Consequently, these findings support the rejection of the null hypothesis, suggesting a significant relationship between age-cohort categorization and perspectives regarding governance technology and policy advocacy.

**Table 3.**
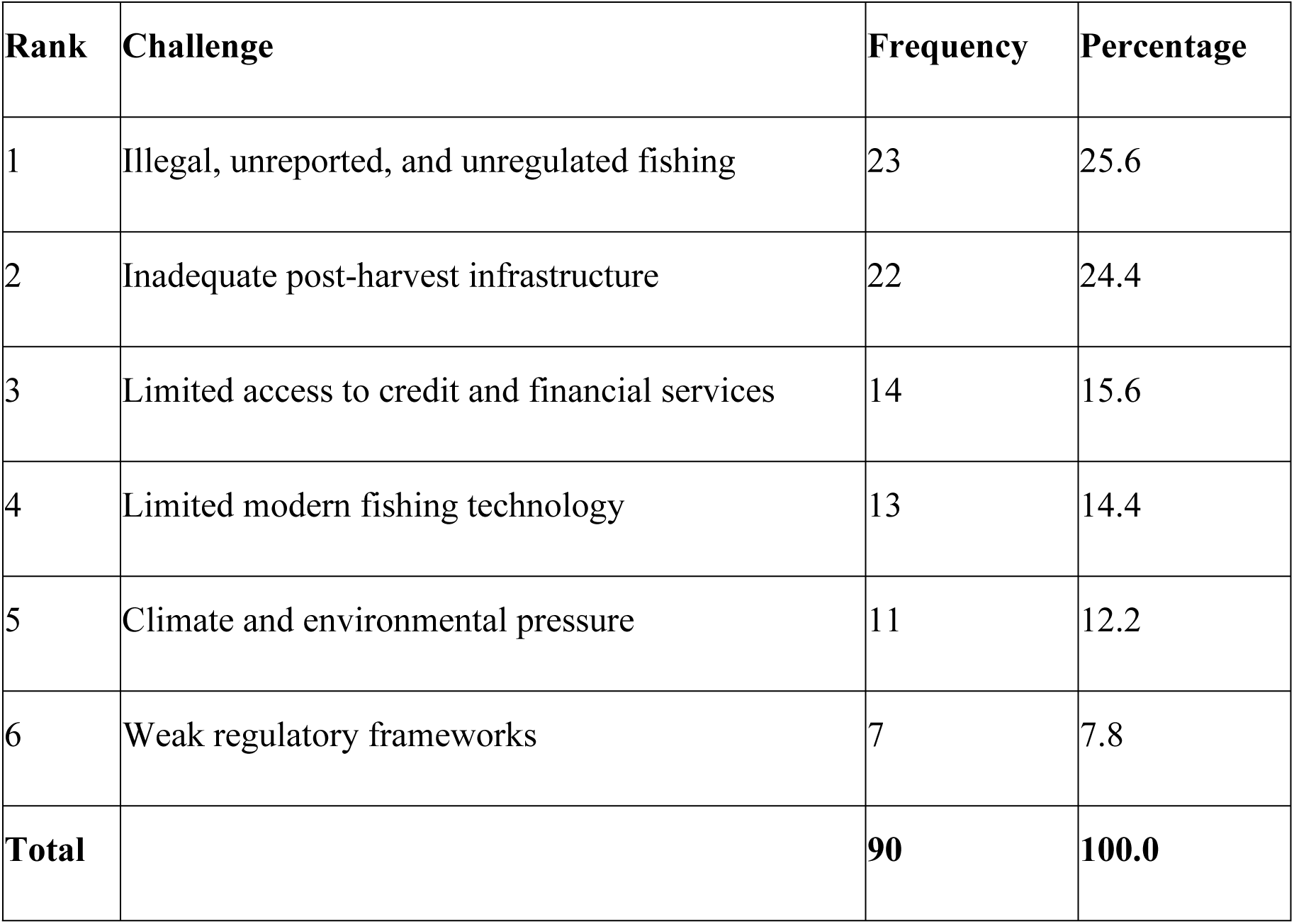
Self-Reported Major Challenges.

Illegal, unreported, and unregulated fishing emerges as the foremost self-identified obstacle, followed closely by deficiencies in infrastructure. Collectively, these two impediments constitute approximately 50% of the total reported constraints. Financial services and technological limitations constitute the intermediate tier, whereas environmental pressures and regulatory weaknesses occupy the tertiary tier. This hierarchy aligns with regional co-management research concerning the Somali Region and neighboring East African small-scale fisheries, where IUU fishing and gaps in cold-chain and post-harvest infrastructure are consistently identified as the primary challenges within fragile-state contexts (Nakamura et al., 2021).

**Figure 1.**
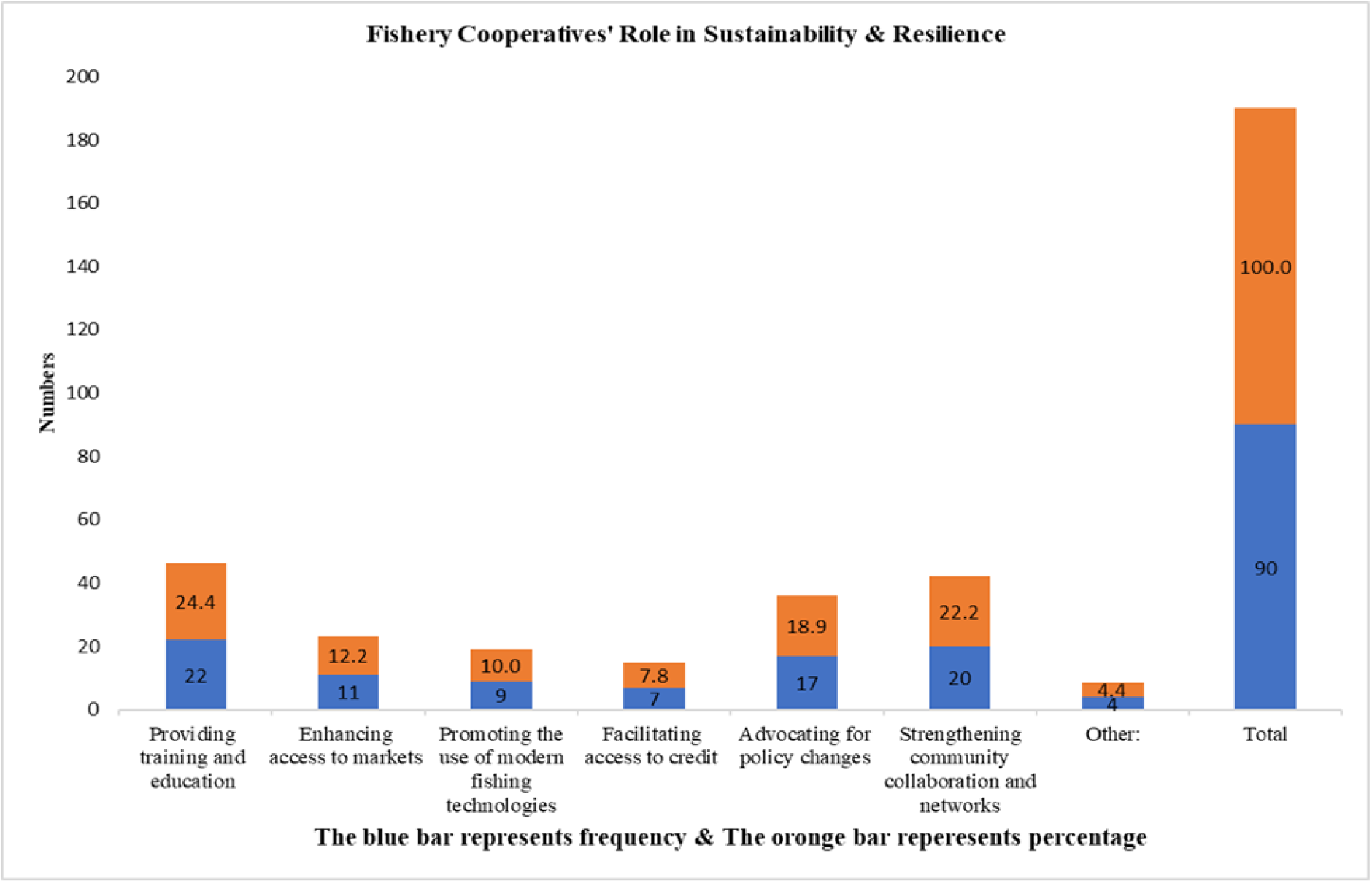
Self-Reported Cooperative Contributions

Training and education, alongside community collaboration, account for nearly half of all reported cooperative contributions. Community collaboration, defined as cooperative-mediated pooled procurement, catch-data exchanges at landing sites, and customary clan-based reciprocal labor arrangements (Murunga et al., 2021), was identified as the second-most prevalent contribution. Policy advocacy and market access constitute a secondary tier, whereas technology promotion, credit facilitation, and other initiatives form the tertiary tier. This rank ordering aligns with broader cooperative fisheries scholarship; for example, recommendations from Ghana suggest that policy interventions should prioritize cooperative capacity building in financial management and governance (Asante, 2024), while evidence from the Mexican lobster sector indicates that strong collective-action characteristics are quantitatively associated with sustainable outcomes (Elsler et al., 2022). These findings corroborate the Social-Ecological Systems framework, establishing governance systems and actor-community dynamics as the primary mechanisms through which training and capacity building enhance cooperative resilience (McGinnis & Остром, 2014).

## 5. Discussion

This discussion is organized into three subsections: an examination of the socio-ecological constraints facing Somali fishery cooperatives, wherein the rank ordering observed in the results is corroborated by international and local institutional evidence; an assessment of cooperative contributions to sustainability and resilience, triangulated with institutional endorsements; and a concluding review of theoretical implications, study strengths, and limitations.

### 5.1 Socio-Ecological Constraints on Cooperative Effectiveness

Illegal, unreported, and unregulated fishing stands as the primary impediment to cooperative efficacy, identified by one-quarter of key informants as the most significant barrier. This assessment is supported by corroborating evidence from international institutional bodies. Research by the Secure Fisheries program, utilizing Automatic Identification System data to monitor foreign fleet operations, estimates that IUU fishing activities within Somali waters involve approximately 300,000–400,000 metric tons annually, a volume that considerably surpasses the sustainable yield of local marine resources (Glaser et al., 2019). Furthermore, the United Nations Special Rapporteur on the Right to Food has highlighted the critical necessity of mitigating illegal fishing to safeguard food security in fragile states (Witbooi et al., 2020). This perspective is integrated into the Somali Federal Ministry of Fisheries and Blue Economy’s mandate, which centralizes the “sustainable development of the fisheries sector” within the federal Blue Economy Framework (Aded, 2023). Additionally, the estimated US$300 million in annual economic losses attributed to foreign-driven IUU fishing aligns with reconstructions by Secure Fisheries, thereby establishing the fundamental rationale for the cooperative co-management model adopted in this research (Omar et al., 2019).

Inadequate post-harvest infrastructure serves as the secondary impediment, with approximately one-quarter of respondents highlighting critical deficits in cold-chain capacity. This challenge is contextualized by World Bank data, which characterizes Somalia’s domestic fisheries as predominantly artisanal, reliant on small-scale vessels utilizing gillnets, handlines, and traps—gear types that account for over 60% of global small-scale fisheries yields (Cashion et al., 2018). Research concerning sub-Saharan Africa indicates that post-harvest quality degradation can account for up to 70% of total losses within fragile-state supply chains (Mramba & Mkude, 2022). Locally, evidence from Puntland suggests that formal cooperative organization can partially mitigate these infrastructure deficits, as organized fishers achieve superior operational outcomes compared to their unorganized counterparts (ABDIRAHMAN, 2022).

Limited financial services, technological barriers, environmental stressors, and regulatory inadequacies constitute the lower-tier constraints. Collectively, these factors account for 50% of the reported limitations. This hierarchy aligns with regional scholarship on small-scale fisheries in fragile states; specifically, post-conflict case studies from Kenya, Tanzania, Mozambique, and Liberia presented at the 2019 Somali Fisheries Forum demonstrate that, where national-level enforcement capacity is constrained, IUU containment and post-harvest infrastructure investment emerge as the dominant dual challenges (Roberts et al., 2019). This structural pattern extends to the governance dynamics observed in South-Central Somalia (ABDIRAHMAN, 2022). The Intergovernmental Authority on Development formally validated this constraint profile during its national assessment of the Blue Economy’s contribution to Somalia’s sustainable development (Mohamed, 2022). Consequently, the Somali Federal Government and Federal Member States have institutionalized a joint revenue-sharing mechanism for fisheries and natural resources, operationalized through the Somali Fisheries Development Framework 2018–2020 of the Eighth National Development Plan (Mohamed, 2022). These instruments operate within the international legal framework provided by the Voluntary Guidelines for Securing Sustainable Small-Scale Fisheries in the Context of Food Security and Poverty Eradication (Aded, 2023).

### 5.2 Cooperative Contributions to Sustainability and Resilience

*The four primary contribution pathways—training and education, community collaboration, conflict resolution, and price negotiation/market development—find their closest international analogue in the literature on Mexican lobster cooperatives. Research indicates that robust collective action—defined empirically by adherence to collective-choice agreements, active monitoring, conflict-resolution mechanisms, and engagement in nested organizational hierarchies—facilitates sustainable fisheries and is quantitatively associated with capturing trade benefits* (Elsler et al., 2022). This empirical pattern aligns with the SES framework’s first-tier variables concerning actor-community and governance systems, providing a validated SES-level finding for Somalia’s hybrid cooperative model, where federal co-management mandates overlap with customary clan-based reciprocal arrangements (McGinnis & Остром, 2014). The Kenyan Beach Management Unit literature corroborates the significance of leadership-and-trust mechanisms (Murunga et al., 2021), while Jamaica’s small-scale fisheries research confirms that geographic proximity and gear-based homophily serve as the operational referents for “community collaboration” (Alexander et al., 2018). These mechanisms and market access strategies receive joint endorsement from international and local institutions. FAO evidence regarding the role of fisheries in food security—citing that fish provide 15% of global animal protein intake and support approximately 600 million livelihoods (Witbooi et al., 2020)—reinforces the relative priority Somali cooperatives assign to training and community collaboration, particularly as small-scale fisheries represent the primary protein source in coastal African diets (Fakoya et al., 2024).

The Federal Ministry of Fisheries and Blue Economy formalizes this emphasis, positioning training and capacity-building as central to cooperative resilience within its vision statement and the NDP8 Somali Fisheries Development Framework (Mohamed, 2022). Regarding policy advocacy, the convergence between the Federal Ministry’s co-management arrangements and the Voluntary Guidelines for Small-Scale Fisheries (SSF) supports the conclusion that cooperative policy engagement is a precondition for sustainable resource management (Nakamura et al., 2021).

Concerning market access, Secure Fisheries’ assessment of regional market geography—specifically Ethiopia and Kenya as nearby, densely populated export destinations—establishes the regional context, while local evidence from the Mogadishu market highlights the structural deficiencies that cooperatives are uniquely positioned to mitigate (Glaser et al., 2019; Perry & Aden, 2023). Conversely, the promotion of modern technology and credit facilitation is categorized within the secondary tier; these are consistent with well-documented international SSF-gap priorities and constitute the primary financial-inclusion challenges identified in the SSF literature (Elegbede et al., 2023).

Ultimately, these five contribution pathways substantiate the theoretical assertion presented in Section 2: Somali fishery cooperatives in South-Central Somalia function as institutional intermediaries analogous to GS2 non-governmental organizations within the SES framework (McGinnis & Остром, 2014). They operationalize the four first-tier SES components—resource systems, resource units, governance systems, and actor communities (ABDIRAHMAN, 2022)—within coastal settlements where, following the post-1991 collapse of centralized fisheries administration and the subsequent, intermittent restoration of federal co-management mandates, national enforcement capacity remains limited (Glaser et al., 2019).

### 5.3 Theoretical implications

The study advances the Social-Ecological Systems framework in two primary respects. First, the data illustrate how Somali cooperatives sustain governance mechanisms notwithstanding the absence of Ostrom’s initial design principle, relying instead on customary clan-based arrangements for compensation. Second, the chi-square analysis revealing an association between age cohorts and support for policy advocacy and modern technology expands the actor-community dimension, suggesting that age-cohort analysis should be adopted as a standard variable in SES assessments within post-conflict fisheries. These findings transcend the conventional substitution–integration dichotomy found in co-management literature (Roberts et al., 2019a, 2019b). Instead, Somali fishery cooperatives emerge as a distinct analytical entity—“hybrid intermediation”—wherein they leverage customary clan governance to address deficits in federal capacity (ABDIRAHMAN, 2022), while functioning concurrently as implementation vehicles for the Federal Blue Economy Framework’s co-management provisions in areas of limited federal penetration (Nakamura et al., 2021). This duality is congruent with the Voluntary Guidelines for SSF, which recognize that cooperatives in fragile environments often operate at the nexus of customary self-organization and state-sanctioned co-management (Aded, 2023). Future scholarly inquiry should determine whether this hybridity constitutes a transitional phase or a stable equilibrium amidst persistent fragility.

## Conclusion

*This study demonstrates that Somali fishery cooperatives function as institutional intermediaries, analogous to GS2 non-governmental organizations within the Social-Ecological Systems framework* (McGinnis & Остром, 2014). By synthesizing the four core SES tiers—resource systems, resource units, governance systems, and actor communities (ABDIRAHMAN, 2022)—these entities operationalize the broader global fisheries contribution into localized practice within the coastal settlements of South-Central Somalia.

Globally, this sector sustains approximately 660–820 million livelihoods, accounting for 10–12% of the global population, and provides roughly 15% of total animal-protein intake (“The State of World Fisheries and Aquaculture 2020,” 2020).

At the cooperative level, intermediation is primarily driven by training, education, community collaboration, policy advocacy, and market access, which collectively constitute 77.7% of reported contributions. This hierarchical structure is consistent with broader literature on African cooperative fisheries, which underscores the necessity of capacity-building in financial management and transparent governance (Asante, 2024). Furthermore, these findings parallel evidence from Mexican lobster fisheries, where robust collective-action mechanisms are quantitatively linked to sustained cooperative performance (Elsler et al., 2022).

Notwithstanding these contributions, cooperatives in South-Central Somalia encounter six primary constraints: Illegal, Unreported, and Unregulated fishing; inadequate post-harvest infrastructure; limited credit; insufficient modern technology; environmental and climate pressures; and regulatory deficiencies (Roberts et al., 2019). The first two factors represent 50.0% of reported limitations and contribute to estimated annual losses of approximately US$300 million—a burden amplified by regional insecurity and systemic regulatory gaps. To mitigate these impediments, this study proposes three interventions to be facilitated via a tripartite partnership involving federal authorities, NGOs, and the private sector (Mohamed, 2022):

➢ Implementation of co-management agreements granting cooperatives exclusive rights in exchange for catch-data reporting, administered by the Federal Ministry of Fisheries and Cooperative Boards.
➢ Creation of revolving credit funds within cooperatives to decrease reliance on informal lenders, led by the Central Bank and international NGOs.
➢ Deployment of solar-powered cold storage at cooperative landing sites via public-private partnerships involving local cooperatives and private sector entities.

These interventions aim to operationalize the Federal Ministry’s Blue Economy Framework and the NDP8 Somali Fisheries Development Framework (Mohamed, 2022), while aligning with the FAO Voluntary Guidelines for SSF (Nakamura et al., 2021) to target critical fishery-cooperative functions. A primary limitation of this study is the gender-imbalanced sample, which deviates from the 74.4% male and 25.6% female composition documented in recent comparable surveys of Mogadishu fisheries (Hassan & Hossain, 2023). Future research should emphasize female participation within post-harvest and retail sectors, as existing literature on Mogadishu small-scale fisheries indicates that women play vital roles in processing, marketing, retailing, and gleaning—activities essential to the resilience of fishery households and local economies (Mohamed et al., 2026). Moreover, systematic reviews have identified these value-chain segments as areas where fisheries research has historically exhibited significant gender bias (Smith & Basurto, 2019).

## Acknowledgments

The authors extend their sincere appreciation to the 90 key informants in Benadir, Hirshabelle, Southwest, Galmudug, and Jubaland for their time, candor, and willingness to share insights during the fieldwork process. We further acknowledge the leadership of the following cooperatives—Alla-Aamin, Dan-kulmis, Tawasal, Wadani, Malable, Rajo, Dayah, Chilani, Danwadaag, Al-Adalla, Garsoor, Hibo, Koyoma, Sabriin Jamiil, and Baqdaad—for their instrumental role in facilitating access to respondents. Additionally, the corresponding author expresses gratitude to the peer reviewers, whose constructive feedback materially improved the rigor and clarity of this manuscript.

## Limitation

The study’s gender distribution—comprising 83.3% male and 16.7% female participants—broadly aligns with FAO benchmarks, which estimate female involvement at 14% for direct capture and 50% for onshore post-harvest activities (Mohamed et al., 2026). This composition may, however, constrain the depth of insight regarding the fish-processing and retail sectors, where women typically play a more prominent role. Consequently, future research should prioritize these downstream segments to more fully elucidate the essential contributions of women to the blue economy.

## Ethics statement

Ethical approval was granted by the Institutional Review Board of Salaam University. Participants were fully informed of the study’s scope and purpose and provided written or verbal informed consent. Participation was anonymous, with no personally identifying information disclosed. The research adhered to the Declaration of Helsinki and Somali national research ethics guidelines.

## Data availability statement

The original contributions presented in this study are included within this article. To ensure participant confidentiality, primary qualitative data from interviews with cooperative members, leaders, and authorities across the five South-Central Federal Member States are restricted. Researchers interested in the cooperative governance and community-led enforcement methodology applied here are referred to the protocols documented by (ABDIRAHMAN, 2022)

## Declaration of Generative AI and AI-assisted technologies

The authors utilized Jenni AI to enhance language clarity and manage bibliographic resources. The authors rigorously reviewed and edited the manuscript, assuming full responsibility for the content of the published article.

